# Detecting visual deficits in retinal degeneration mice using photoacoustic tomography

**DOI:** 10.64898/2026.01.30.702640

**Authors:** Guan Xu

## Abstract

We established a photoacoustic tomography and ultrasound imaging system capable of resolving visually evoked hemodynamic responses in the cortical and subcortical visual regions of the brains of freely behaving mice. By searching for anatomical landmarks in the US imaging planes, we can locate brain regions of interest and continuously record HR in these regions. The system was examined in retinal degeneration mice using a 60-second protocol and in Claudin 5 knockdown mice and mice with vision deficits using a 100-minute-long vision research protocol. We found that: 1) visually evoked HR amplitudes increase as visual stimulation intensity increases in both scotopic and photopic conditions; and 2) HR amplitudes increase during the light adaptation time course.

## 1. Introduction

In the brain, many neural pathways signal through disparate nuclei and cortical areas to analyze different aspects of the internal and external environments and then produce appropriate responses. Some of the best studied neural pathways are in the visual system where RGCs signal to >20 brain nuclei to mediate many functions. Researchers analyze various mouse models’ phenotypes to learn how corresponding human diseases might impact visual functions and to assess therapeutic efficacy [1]. Many techniques exist for screening retinal phenotypes [2] but significant challenges remain for efficiently probing higher visual areas.

Photoacoustic tomography (PAT) has recorded visual-evoked HR in V1 in anesthetized [3, 4] by us. PAT images are naturally co-registered with the US images acquired by the same transducer array. Exploiting the acoustic impedance contrast within the brain and transmission beamforming, US imaging reveals detailed brain anatomies. Our study [4] from an anesthetized wild-type (WT) mouse which represent, to our knowledge, the first successful application of PAT without exogenous contrast agent in any rodent species’ subcortical nuclei. We have also quantified visual impairments in rod/cone-degenerate (*rd1)* mice and melanopsin-knockout (“melko”) mice [4]. This low-cost PAT system is easy for other labs to duplicate and thus will greatly benefit the wider research community.

In this study, we developed a prototype PAT+US headmount that can capture the hemodynamics responses in free-moving mice. The headmount imaging system was validated by the well-established retina degeneration and Claudin 5 knock down models in mice.

## 2. Methods

### 2.1 Animal protocols

All animal protocols in this study have been approved by the Institutional Animal Care & Use Committee (IACUC) at the University of Michigan.

Retina degeneration (rd1) mice in our previous publications [3, 4] were used in a 60-second, photopic protocol. Claudin5 knockdown mice were used in a long protocol (>100 minutes) including both photopic and scotopic phases.

### 2.2 System configuration

As shown in Fig. 1, our system consists of a heamount including all the PAT and US measurements at the mouse brain, a tunable laser for PAT illumination (Opolette, OPOTEK) and an US platform (Vantage 256, Verasonics). The laser emission is synchronized with the US system reception. A frame rate of 10Hz is achieved, which is sufficient to capture hemodynamic responses of visual stimulation that happens in the time scale of second.

**Fig. 1.**
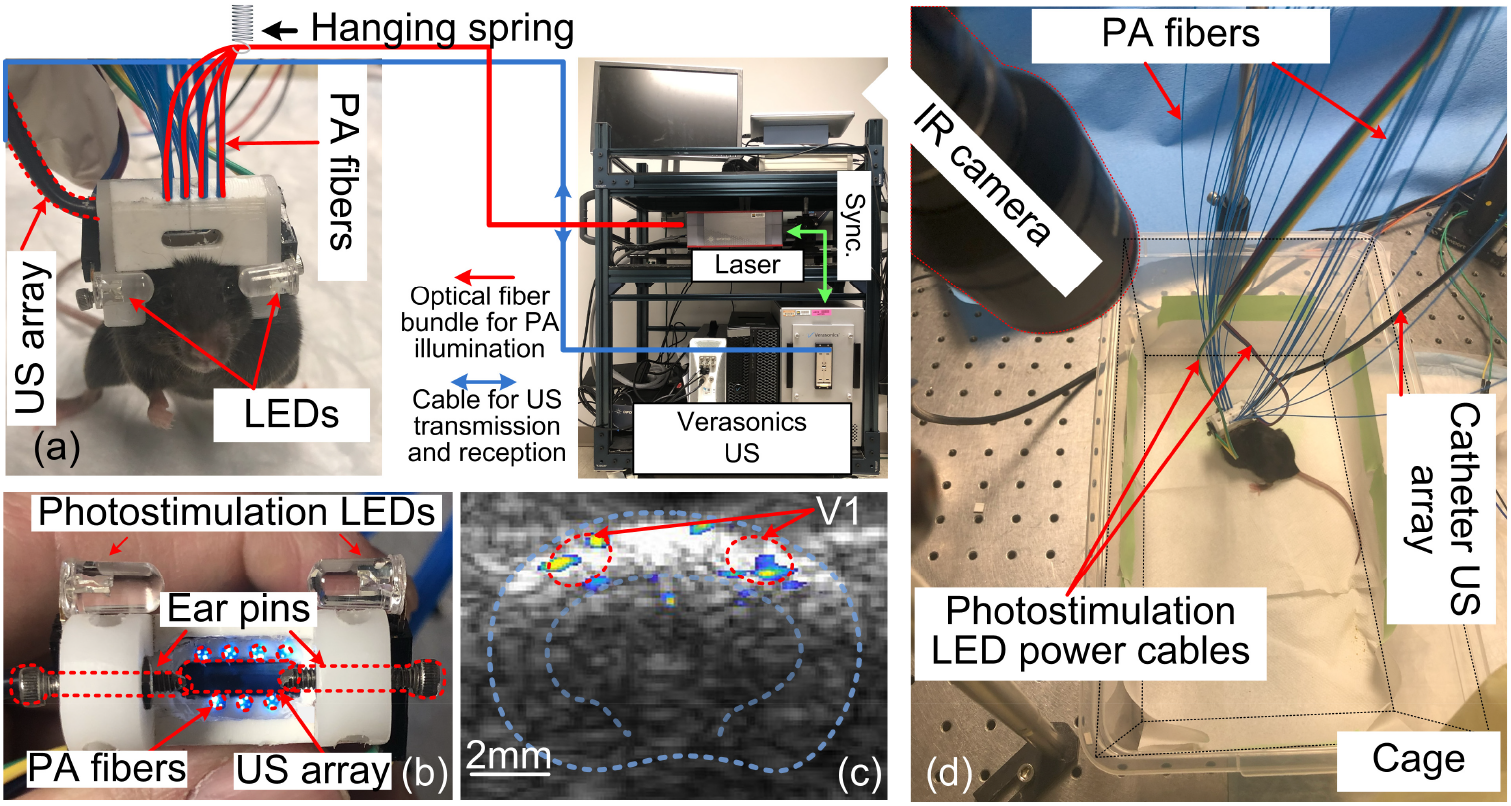
Experiment setup. (a) Mouse with the headmount installed. The data acquisition system is integrated to a mobile cart. (b) Bottom view of the headmount. (c) representative image acquired by the system. *Blue and green contours* mark the outlines of skull surface and inner brain. The functional map showing the brain regions with evoked hemodynamic activities is coregistered. (d) Free-moving mouse experiment configuration. An Infrared (IR) camera captures the mouse’s movements throughout the procedure.

### 2.3 Headmount for mouse brain imaging

We made a prototype head-mount probe containing 28 optical fibers (FT300EMT, Thorlabs) for delivering PA illumination (Fig. 2a,b). A 1D piezoelectric catheter US array (Acunav Siemens) was integrated into the headmount. The array had an 8MHz central frequency, -3 dB bandwidth of 3-10 MHz, and 64 elements at a 0.1mm pitch. The headmount had 2 LEDs to stimulate both retinas. US coupling gel filled the gap between the headmount and the mouse. Two ear pins fixed the headmount to the skull. The headmount was very stable in a >1-hour imaging protocol, to be presented in Aim 3. Analgesia was administered. We located the V1+SC plane by rotating the US catheter to find the largest contour of the inner brain (Fig. 2c *green contour*). The mouse was put in a 20cm (W) × 40cm (L) × 15cm (D) cage. A spring posi-tioned 3 ft above the cage counter-weighted the headmount and cable/fibers within ±4g and the cable/fibers were loose enough (Fig. 2d) to let the mouse move freely. The cage was illuminated by invisible infrared LEDs, and a camera recorded mouse activity (Fig.2d). To avoid locomotion-induced modulation of vHR, we will only include data from imaging trials lacking locomotion.

**Fig. 2.**
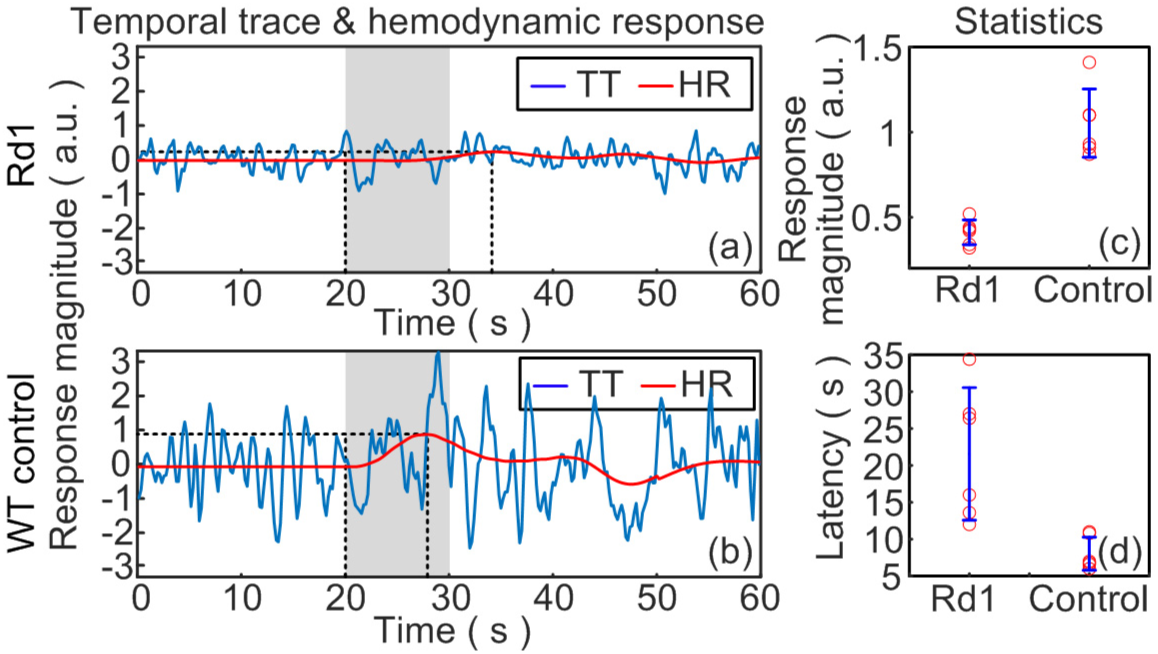
Results from rd1 mice. (a-b) Average temporal trace (TT) and visually evoked hemodynamic responses (HR) extracted from rd1 (n=6) and control mice (n=6), respectively. Shaded areas are the 10-second time period of visual stimulations. (c-d) Statistics of the HR response magnitudes and latency derived from 6 rd1 and 6 control mice.

### 2.4 Data processing

We established a data processing protocol to quantify the visually-evoked hemodynamics responses (vHR) in our recent study [5]. In brief, we filter out repetitive noises caused by respiration, heartbeat, low-frequency baseline signals, and autocorrelated measurement noises. These operations remove uncertainties, including optical attenuation in individual animals, and retain vHR. With limited pilot data, we used the fixed canonical shape HR function [6-8], an empirical model frequently used in fMRI [9] and fNIRs [10, 11], to model the impulse HR to neuronal activation. The impulse response (*h*_*t*_) was convolved with our photostimulation sequence (*S*_*t*_) forming the basis function (*x*_*t*_). Following standard methods in [12-17], the vHR was modeled as the linear combination of a basis set, as the summation of a constant bias, the second-order Taylor expansion of the basis function, and a noise term. Since some mouse strains with visual deficits have delayed responses, we fit the temporal traces at each pixel in PA image to a series of basis sets with 1-sec shifts. The basis set with least fitting error and lowest *p*-value in linear fit was chosen as the isolated vHR.

## 3. Results

The prototype system can detect different vHR in V1 in rd1 and wild-type control mice with their averaged responses being, respectively, ∼90% (*p*<1×10^−4^) and ∼50% (*p*>0.05) higher-amplitude in free-moving mice (Fig. 2). These results show that anesthesia-induced weaker vHR are harder to detect reliably and to quantify accurately, reinforcing the need to develop PAT for unanesthetized mice.

Fig. 2(a-b) show average vHR in free-moving rd1 and wild-type control mice. In Fig. 2(c-d), the rd1 mice show lower response magnitude (p<0.001) and longer latency (p<0.02) .

We also tested a wider range of visual stimuli and ambient lighting conditions. We used the prototype probe to image V1 in free-moving Cldn5KD mice, following the 62-min protocol in Fig. 3 *top* to measure vHR under dark-adapted state, then turned on a constant background light to track the progression of light adaptation, and finally under fully light-adapted state. Pilot data from 2 Cldn5KD mice and 1 control not only found that Cldn5 deficiency impairs image-forming vision but also yielded additional insights: *1*) both dark- and light-adapted vHR are weakened (Fig. 3b *left & right plots*), suggesting deficits in rod-as well as cone/melanopsin-mediated signaling; *2*) light adaptation seems to take longer (Fig. 3b *middle plot*), implying deficits in light-adaptive mechanisms e.g. dopaminergic neuromodulation ; *3*) HR latencies remain WT-like (not shown), so rods/cones rather than the much slower melanopsin remain the primary contributors to image-forming vision.

**Fig. 3.**
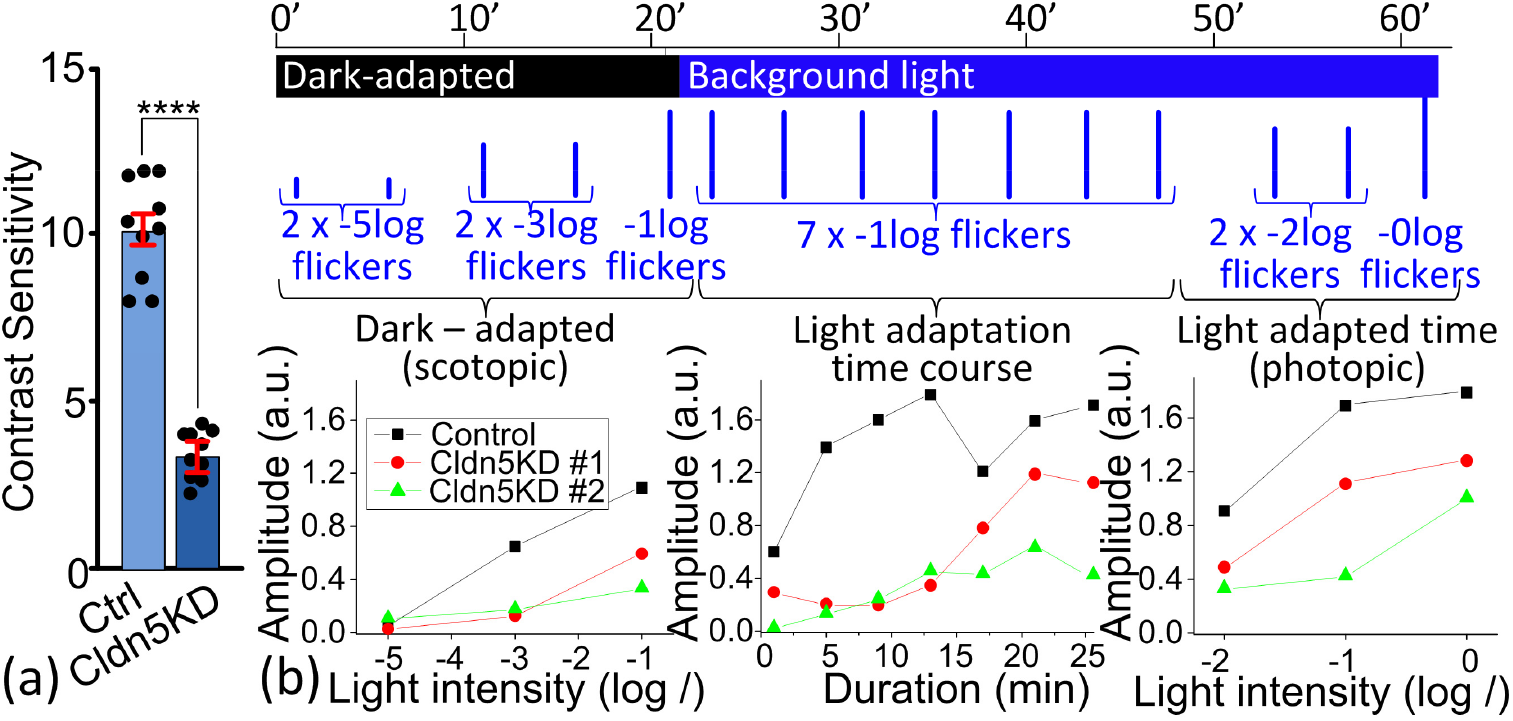
Knocking down Cldn5 impairs visual functions. (a) Optomotor data. ****, *p*<0.0001. (b) Visual stimulation protocol for PAT (*top diagram*) and V1 data from free-moving mice (*bottom plots*). All flickers are 10× 1Hz 500ms flashes. The -5, -3 and -2 log intensities are given twice to allow vHR averaging. We use the first 5 vHR to plot dark-adapted vHR amplitudes (*left plot*), the next 7 HR to track amplitude increase during light adaptation (*middle plot)*, and the final 4 vHR to plot light-adapted amplitudes (*right plot*).

## 4. Discussion

The central frequency of the miniaturized US transducer, 8MHz, in the free-moving mice setup is lower than the 10-MHz central frequency US transducer in our previous studies with anesthetized mice. This leads to slightly lower resolution in the images. However, we are focusing on examining the feasibility of the proposed coronal imaging plane headmount in this study. In the future study, we will integrate US transducers with a broader bandwidth for higher spatial resolution and sufficient penetration.

The protocol for rd1 mice online includes a photopic phase. The temporal traces show a slightly low signal-to-noise ratio compared to our previous studies with a scoptopic setup. However, our data processing methods successfully extracted the visually evoked HR that agrees with the phenotypes of the established mouse models.

The headmount design in this study shows reliability in capturing HR in mouse brain in extended protocols. In the future works, we will further reduce the volume and weight of the headmount.

## 5. Conclusion

We developed a prototype photoacoustic ultrasound imaging system for capturing visually evoked hemodynamic responses in the mouse brain. The system was validated by established mouse models. We will implement the system in a broader range of applications in the future.

## References

1. J. Won, L. Y. Shi, W. Hicks, J. Wang, R. Hurd, J. K. Naggert, B. Chang, and P. M. Nishina, “Mouse Model Resources for Vision Research,” Journal of Ophthalmology 2011, 391384 (2011).

2. J. Won, L. Y. Shi, W. Hicks, J. Wang, J. K. Naggert, and P. M. Nishina, “Translational Vision Research Models Program,” in Retinal Degenerative Diseases, (Springer US, 2012), 391–397.

3. K.-W. Chang, Y. Zhu, X. Wang, K. Y. Wong, and G. Xu, “Label-free photoacoustic computed tomography of mouse cortical responses to retinal photostimulation using a pair-wise correlation map,” Biomedical Optics Express 13, 1017–1025 (2022).

4. K.-W. Chang, E. Belekov, X. Wang, K. Y. Wong, Ö. Oralkan, and G. Xu, “Photoacoustic imaging of visually evoked cortical and subcortical hemodynamic activity in mouse brain: feasibility study with piezoelectric and capacitive micromachined ultrasonic transducer (CMUT) arrays,” Biomedical Optics Express 14, 6283–6290 (2023).

5. K.-W. Chang, X. Wang, K. Wong, and G. Xu, “Label-free photoacoustic computed tomography of visually evoked responses in the primary visual cortex and four subcortical retinorecipient nuclei of anesthetized mice,” Neurophotonics 11, 035005 (2024).

6. R. Henson and K. Friston, “Convolution models for fMRI,” Statistical parametric mapping: The analysis of functional brain images, 178–192 (2007).

7. T. Jin and S.-G. Kim, “Cortical layer-dependent dynamic blood oxygenation, cerebral blood flow and cerebral blood volume responses during visual stimulation,” NeuroImage 43, 1–9 (2008).

8. M. Havlicek, D. Ivanov, A. Roebroeck, and K. Uludağ, “Determining Excitatory and Inhibitory Neuronal Activity from Multimodal fMRI Data Using a Generative Hemodynamic Model,” Frontiers in Neuroscience 11(2017).

9. M. J. McKeown, L. K. Hansen, and T. J. Sejnowsk, “Independent component analysis of functional MRI: what is signal and what is noise?,” Current Opinion in Neurobiology 13, 620–629 (2003).

10. E. Hernandez-Martin, F. Marcano, C. Modroño, N. Janssen, and J.L. González-Mora, “Difuse optical tomography to measure functional changes during motor tasks: a motor imagery study,” Biomedical Optics Express 11, 6049–6067 (2020).

11. Z. Cai, A. Machado, R. A. Chowdhury, A. Spilkin, T. Vincent, Ü. Aydin, G. Pellegrino, J.-M. Lina, and C. Grova, “Difuse optical reconstructions of functional near infrared spectroscopy data using maximum entropy on the mean,” Scientific Reports 12, 2316 (2022).

12. K. Han, H. Wen, J. Shi, K.-H. Lu, Y. Zhang, D. Fu, and Z. Liu, “Variational autoencoder: An unsupervised model for encoding and decoding fMRI activity in visual cortex,” NeuroImage 198, 125–136 (2019).

13. F. Cignetti, E. Salvia, J.-L. Anton, M.-H. Grosbras, and C. Assaiante, “Pros and Cons of Using the Informed Basis Set to Account for Hemodynamic Response Variability with Developmental Data,” Frontiers in Neuroscience 10(2016).

14. D. H. Kim, K.-D. Lee, T. C. Bulea, and H.-S. Park, “Increasing motor cortex activation during grasping via novel robotic mirror hand therapy: a pilot fNIRS study,” Journal of NeuroEngineering and Rehabilitation 19, 8 (2022).

15. V. D. Calhoun, M. C. Stevens, G. D. Pearlson, and K. A. Kiehl, “fMRI analysis with the general linear model: removal of latency-induced amplitude bias by incorporation of hemodynamic derivative terms,” NeuroImage 22, 252–257 (2004).

16. H. Lambers, M. Segeroth, F. Albers, L. Wachsmuth, T. M. van Alst, and C. Faber, “A cortical rat hemodynamic response function for improved detection of BOLD activation under common experimental conditions,” NeuroImage 208, 116446 (2020).

17. M. A. Lindquist, J. Meng Loh, L. Y. Atlas, and T. D. Wager, “Modeling the hemodynamic response function in fMRI: Eficiency, bias and mis-modeling,” NeuroImage 45, S187–S198 (2009).

18. H. C. van Diepen, A. Ramkisoensing, S. N. Peirson, R. G. Foster, and J. H. Meijer, “Irradiance encoding in the suprachiasmatic nuclei by rod and cone photoreceptors,” The FASEB Journal 27, 4204–4212 (2013).

19. E. Drouyer, C. Rieux, R. A. Hut, and H. M. Cooper, “Responses of Suprachiasmatic Nucleus Neurons to Light and Dark Adaptation: Relative Contributions of Melanopsin and Rod–Cone Inputs,” The Journal of Neuroscience 27, 9623–9631 (2007).

